# Upregulation of PD-L1 as a putative mechanism of resistance to CD47 inhibition in non-small cell lung cancer

**DOI:** 10.64898/2026.04.24.720733

**Authors:** Asa P. Y. Lau, Kristyna A. Gorospe, Kelsie L. Thu

## Abstract

CD47 is a “don’t eat me” signal that suppresses macrophage-mediated phagocytosis. Its upregulation in lung and other cancers facilitates tumour immune escape, making CD47 a promising immunotherapeutic target. Studies have demonstrated anti-tumour efficacy of CD47 blockade in preclinical lung cancer models, but monoclonal antibodies targeting CD47 have had limited efficacy as monotherapy in solid tumour patients to date. This discrepancy may in part reflect the use of human tumour xenografts in mice that do not have fully-functioning immune systems in preclinical efficacy studies. Thus, understanding tumour responses to CD47 inhibition using immune competent lung cancer models is needed to inform strategies to harness its therapeutic potential. Here, we characterized the effects of CD47 knockout (KO) on tumour growth and immune responses in two syngeneic, orthotopic murine lung cancer models, LLC-Luc (LLC) and CMT167 (CMT). As expected, CD47 KO impaired the fitness of LLC and CMT cells *in vivo*. Mice with CD47-deficient tumours exhibited prolonged survival and increased infiltration of anti-tumour leukocytes. However, although CD47 KO impaired lung tumour growth in syngeneic mice, KO tumours were ultimately lethal. Immunophenotyping revealed an increased prevalence of PD-L1+ cells in CD47-deficient tumours, nominating PD-L1-mediated suppression of tumour immunity as an acquired mechanism of resistance to CD47 blockade. Concordantly, dual inhibition of CD47 and PD-L1 extended the survival of CMT tumour-bearing mice compared to inhibition of either alone. These findings suggest that PD-L1 blockade could be leveraged to overcome resistance and potentiate the efficacy of CD47-targeted immunotherapy in lung cancer.

## 1.0 Introduction

Discovery of mechanisms enabling tumour immune evasion stimulated the development of immunotherapies like PD-1/PD-L1/CTLA-4 blockade that re-invigorate tumour immunity to induce tumour killing.^1^ Although these drugs elicit remarkable tumour regressions and remissions in some non-small cell lung cancer (NSCLC) patients, their clinical utility is restricted to eligible patients, of which only 15-20% respond, and limited by innate and acquired tumour resistance.^1,2^ Thus, additional immunotherapies are needed to benefit NSCLC patients who cannot be treated with approved immune checkpoint inhibitors.

CD47 is a transmembrane protein frequently overexpressed and associated with poor prognosis in NSCLC.^3^ Binding of CD47 to its receptor SIRPɑ, on phagocytes like dendritic cells and macrophages, transmits a potent anti-phagocytic signal that prevents cytoskeletal rearrangements required for these cells to carry out phagocytosis.^4^ Cancer cells hijack the immunosuppressive function of CD47 to escape phagocyte-mediated destruction and its stimulation of tumour immunity. Despite promising data in preclinical NSCLC models, clinical trial results indicate that CD47-targeted immunotherapy, or CD47 blockade, administered using antibody-based therapeutics has limited efficacy in solid tumour patients. This discordance may be attributable, at least in part, to the fact that most preclinical studies have evaluated CD47 blockade in human NSCLC xenografts grown in immunodeficient mice, which do not recapitulate tumour-immune interactions critical for modelling stimulation of anti-tumour immune responses induced by CD47 blockade.^5^ It could also reflect limited knowledge regarding the mechanisms underlying tumour response and resistance to CD47 inhibition, which must be understood to inform strategies for using CD47 blockade effectively in the clinic.^6^

In this study, we sought to evaluate CD47 blockade in immune competent NSCLC models to understand its effects on tumour growth and the tumour microenvironment in syngeneic hosts, and to identify putative mechanisms of resistance. We discovered that CD47 knockout (KO) in NSCLC cells impairs tumour growth in syngeneic mice, but we also found that CD47-deficient tumours are ultimately lethal. We observed that PD-L1+ cells were more prevalent in CD47 KO tumours and that inhibiting PD-L1 in CD47-deficient tumours prolonged mouse survival. Taken together, our findings suggest that PD-L1 confers resistance to CD47 blockade, and consequently, that combining CD47 and PD-L1 blockade could be an effective strategy for potentiating the efficacy of CD47-targeted immunotherapy in NSCLC.

## 2.0 Materials and Methods

### 2.1 NSCLC models

LLC-Luc (LLC) and CMT167 (CMT) cell lines were obtained from Promega and Sigma, respectively, and cultured according to supplier instructions. Knockdowns of CD47 and CD274 (the gene encoding PD-L1) were achieved using lentivirally delivered CRISPR constructs (Cas9 and sgRNA) and clonal knockout lines were established as previously described.^7^ sgRNA sequences are provided in **Supplementary Table 1**. Cell lines were confirmed to be mycoplasma negative using a PCR-based assay.

### 2.2 Multi-colour competition assays (MCAs)

LLC and CMT cells with stable Cas9 expression were transduced with mCherry- or GFP-tagged pLentiguide vectors to express sgRNA targeting LacZ (sgLacZ, negative control) or GFP-tagged lentivectors to express sgRNA targeting CD47 (sgCD47-1, sgCD47-2). mCherry+ sgLacZ cells were mixed with GFP+ sgLacZ or GFP+ sgCD47 cells at a 1:1 ratio and orthotopically implanted into the left lung of 6-8 week old C57BL/6 mice, as previously described.^8^ The abundance of GFP+ and mCherry+ cells in cell mixtures prior to implantation and in dissociated tumours harvested at humane endpoints was quantified using flow cytometry (described below).

### 2.3 Animal studies

Animal studies were performed following an institutionally approved protocol in accordance with guidelines of the Canadian Council on Animal Care, which are compliant with ARRIVE guidelines. For orthotopic tumour models, CMT (5 x 10^5^) or LLC (2 x 10^5^) cells were injected into the left lung of 6-8 week old female or male syngeneic C57BL/6 mice, respectively. Bioluminescence imaging (BLI) of LLC tumours expressing luciferase was done on a Newton 7.0 FT-500 Bioluminescent Animal Imager (Scintica) following intraperitoneal (IP) injection of 2.25mg luciferin every 2-3 days to monitor orthotopic LLC tumour growth in real-time. Mice with orthotopic tumours were euthanized at humane endpoints associated with lung tumour burden (eg. hunched posture, failure to groom, difficulty breathing) and survival was analyzed using Kaplan-Meier survival curves.

For therapeutic studies, LLC and CMT cells were orthotopically injected as described above. CMT tumours were also modelled subcutaneously by injecting 4 x 10^5^ cells into the right hind flank of 6-8 week old syngeneic female C57BL/6 mice. The efficacy of CD47-targeted monoclonal antibodies (mAb) was tested against orthotopic LLC tumours and subcutaneous CMT tumours. Established LLC tumours were treated with 200µg of anti-CD47 mAb (clone miap301) or IgG2a isotype control (2A3) administered IP on Days 4-9. Tumour growth was monitored with BLI imaging. Subcutaneous CMT tumours were treated with 100µg anti-CD47 or isotype control dosed intratumourally every 3 days for 5 total doses and tumour growth was monitored with calipers. Anti-tumour effects of PD-1 blockade (clone RMP1-14 or IgG2a isotype control) were tested against orthotopic LLC tumours (150µg IP on Days 3, 6, 9) with BLI done to monitor tumour growth. The efficacy of PD-L1 blockade (clone 10F.9G2 or IgG2b isotype control) was tested on subcutaneous CMT tumours treated with intratumoural injections of 100µg every 3 days for 5 total doses, and tumour growth was monitored with calipers.

To mimic combined treatment with CD47 and PD-L1 blockade, we conducted genetic studies to determine how dual CD47 and PD-L1 loss of function (LOF) affected lung tumour progression. WT and CD47 KO LLC and CMT cells with stable Cas9 expression were transduced with sgLacZ or sgRNA targeting CD274 (the gene encoding PD-L1; **Supplementary Table 1**). These cells were orthotopically injected into C57BL/6 mice as above, and mice were euthanized at humane endpoints. We also evaluated combined CD47 and PD-L1 blockade using a pharmacologic approach. WT and CD47-deficient CMT cells were orthotopically injected into syngeneic mice, as above, since CD47-targeted antibodies ineffectively suppressed tumour growth in mice (see results below). On Day 3, mice in single agent and upfront combination treatment groups were administered 200µg of anti-PD-L1 mAb or IgG2b isotype control by IP injection every 3 days for 4 doses total. For the sequential treatment group, mice were treated with the same number of anti-PD-L1 or isotype control antibody doses starting at Day 15. Anti-tumour efficacy was evaluated by comparing Kaplan-Meier survival curves.

### 2.4 Flow cytometry and immune phenotyping

Flow cytometry was used to measure CD47 and PD-L1 expression on LLC and CMT cells lifted with TrypLE. To confirm loss of PD-L1 expression in CRISPR edited lines, cells were treated with 100ng/ml IFNγ for 24hr. For immune phenotyping studies, resected tumours were dissociated into single cell suspensions using mouse tumour dissociation kits with physical disruption using a GentleMACs dissociator (Miltenyi). Following dissociation, red blood cells were lysed. Cells were stained with antibodies and DAPI or Zombie Violet as a viability marker, and analyzed on a Sony SP6800 Spectral Cytometer or Cytoflex-LX cytometer. Tumours were harvested prior to humane endpoints, with WT tumours harvested on Day 6 (LLC) and Day 10 (CMT), and KO tumours harvested on Day 15 for both models. These time points were estimated to represent the midpoint of the tumour growth studies. Antibodies used for flow cytometry and immune profiling are listed in **Supplementary Table 2**. Cell populations were analyzed in FlowJo using gating strategies shown in **Fig. S1**.

### 2.5 Statistical analysis

Statistical tests were performed using GraphPad Prism V10 and are specified in the figure legends. Continuous data were analyzed with unpaired, two-tailed student’s t-tests. Log-rank tests were used to analyze Kaplan Meier Survival curves. RNA expression data for *CD47* and *CD274* in 510 lung adenocarcinoma tumours from the The Cancer Genome Atlas (TCGA) cohort was accessed and analyzed for mutual exclusivity using cBioPortal.^9–11^ For all tests, p < 0.05 was considered significant with *p < 0.05, **p < 0.01, ***p < 0.001, ****p < 0.0001, and ns = non-significant.

## 3.0 Results

### 3.1 CD47 deficiency reduces NSCLC fitness and delays tumour growth, prolonging the survival of immune proficient tumour-bearing mice

We previously showed that CD47 KO has no effect on NSCLC cell fitness or proliferation *in vitro*.^*7*^ To determine how CD47 influences NSCLC fitness and tumour growth in syngeneic mice, we compared the fitness of *Kras*-mutant NSCLC cells (LLC and CMT) engineered with and without CD47 knockdown (sgCD47) **(Fig 1A)** using *in vivo* multicolor competition assays. Briefly, GFP+ sgCD47 cells were mixed 1:1 with mCherry+ or GFP+ control (sgLacZ) cells and cell mixtures were orthotopically injected into immune competent, syngeneic mice. Flow cytometry to quantify the abundance of GFP+ and mCherry+ cancer cells in tumours collected at endpoint indicated a depletion of GFP+ sgCD47 cells relative to mCherry+ sgLacZ cells in both tumour models **(Fig 1B-C)**, indicating that NSCLC cells with CD47 LOF have a fitness disadvantage *in vivo*.

**Figure 1.**
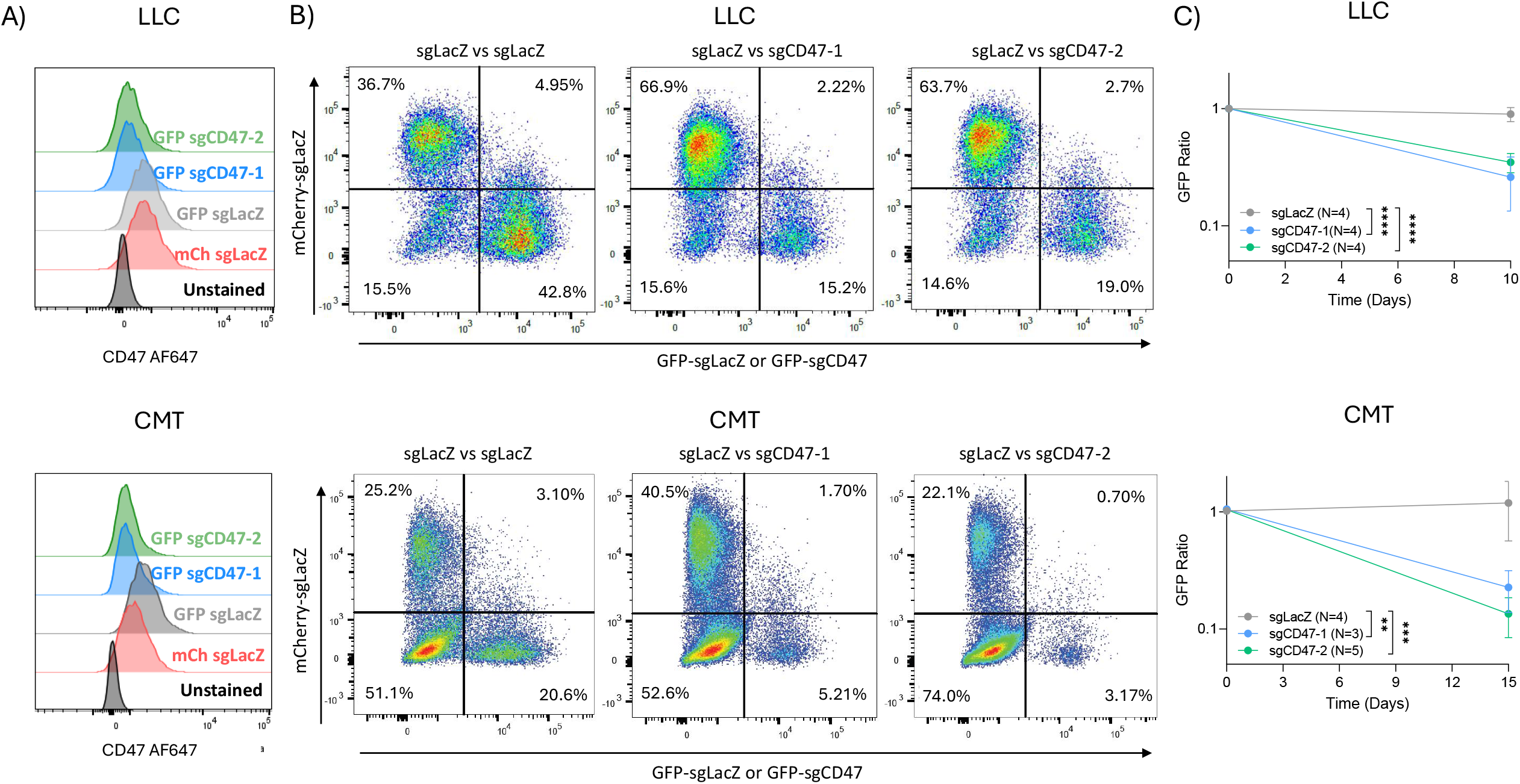
CD47 loss of function reduces tumour cell fitness in syngeneic mice. **A)** Flow cytometry verifying loss of CD47 surface expression in cell lines for competition assays prior to implantation in mice. **B)** Representative flow cytometry plots showing mCherry and GFP populations in competition assay tumours harvested from mice at humane endpoints. The percentage of cells in each quadrant is shown for tumours comprised of each cell mixture (mCherry+sgLacZ vs GFP+sgLacZ; mCherry+sgLacZ vs GFP+sgCD47-1; mCherry+sgLacZ vs GFP+sgCD47-2). **C)** Ratios of GFP+ to mCherry+ cells in competition assay tumours. GFP+ ratios in tumours at endpoint were normalized to GFP+ ratios in cell mixtures prior to orthotopic injection. For CMT, N=4 for sgLacZ vs sgLacZ; N=3 for sgLacZ vs sgCD47-1; N=5 for sgLacZ vs sgCD47-2. For LLC, N=4 for each cell mixture. Error bars indicate mean ± standard deviation. Asterisks indicate significance for unpaired, two-tailed t-tests comparing GFP+ ratios across genotypes.

To investigate the influence of CD47 on tumour growth and progression, we leveraged LLC and CMT lines with CD47 knockout (KO) **(Fig 2A,C**). We observed that syngeneic mice with orthotopic CD47 KO tumours exhibited significantly prolonged survival compared to mice with wildtype (WT) tumours **(Fig 2B,D)**. For the LLC model, which stably expresses luciferase for monitoring tumour growth *in vivo*, the survival benefit seen in KO tumour-bearing mice was associated with slower tumour growth **(Fig S2A-B)**. These studies indicate that CD47 promotes tumour growth and progression in immune competent mice, confirming its therapeutic potential in NSCLC.

**Figure 2.**
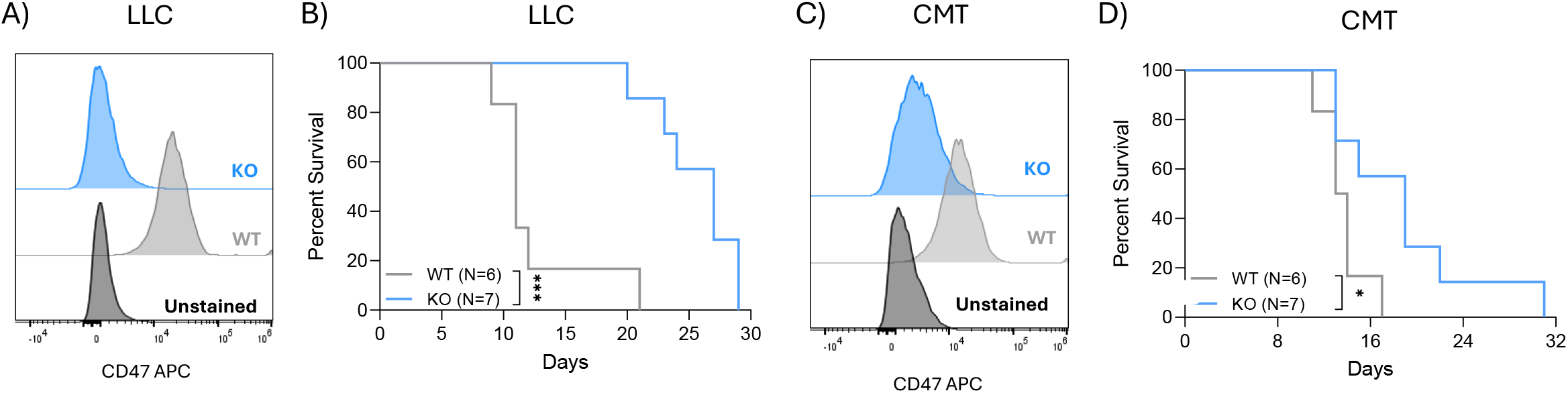
CD47 loss of function (LOF) prolongs survival in immune competent hosts. **A,C)** Flow cytometry confirming loss of CD47 protein expression in LLC and CMT CD47 KO lines. **B**,**D)** Kaplan-Meier survival curves for mice orthotopically injected with LLC and CMT WT (wildtype) or CD47 KO cells (N=6 WT and N=7 KO for each model). Asterisks indicate significance for comparisons of survival between WT versus KO tumour-bearing mice using a log-rank test.

### 3.2 CD47 KO tumours have increased infiltration of anti-tumour immune cells and PD-L1+ cells

We next characterized how CD47 KO affects tumour immune contexture by immune phenotyping of a second cohort of tumour-bearing syngeneic mice (**Fig S1**). Tumours were harvested prior to humane endpoints, at the approximate midpoint of our tumour growth studies **(Fig 2B,D; Fig 3A-B)**. Flow cytometry analyses of tumour immune infiltrates revealed a significant increase in anti-tumour M1-like macrophages (CD64+MHC-II^hi^CD80+) in CD47 KO compared to WT tumours in both models **(Fig 3C-D)**, consistent with enhanced innate immune responses upon CD47 blockade. LLC and CMT tumours displayed differential infiltration of CD8+ T cells, with a reduction of CD8+ T cells evident in LLC KO tumours and an increase in CMT KO tumours **(Fig 3C-D)**. No significant differences in CD4+ T cell infiltration were observed between WT and CD47 KO tumours **(Fig S3)**. LLC KO tumours exhibited higher CD4+/CD8+ T cell ratios compared to WT tumours, while the opposite was seen in CMT KO tumours **(Fig S3)**. Although they had fewer infiltrating CD8+ T cells, CD47-deficient LLC tumours had an increased abundance of CD69+ activated CD8+ T cells, while a decrease in CD69+ CD8+ T cells was evident in CMT KO tumours **(Fig 3C-D)**. The differing phenotypes associated with CD47 KO in LLC and CMT tumours likely reflect inherent differences in their immunosuppressive capabilities and how they stimulate tumour immunity, which has previously been described with respect to their responses to PD-1/PD-L1 blockade and IL-9 therapy.^12–14^ Despite these differences, we observed trending or significant increases in the abundance of CD45-PD-L1+ cells (including tumour cells) and significant increases in CD11b+ PD-L1+ (myeloid) cells in both LLC and CMT KO tumours **(Fig 3C-D)**, suggesting that KO tumours may recruit PD-L1+ cells or upregulate its expression to enable immune evasion when challenged with CD47 blockade.

**Figure 3.**
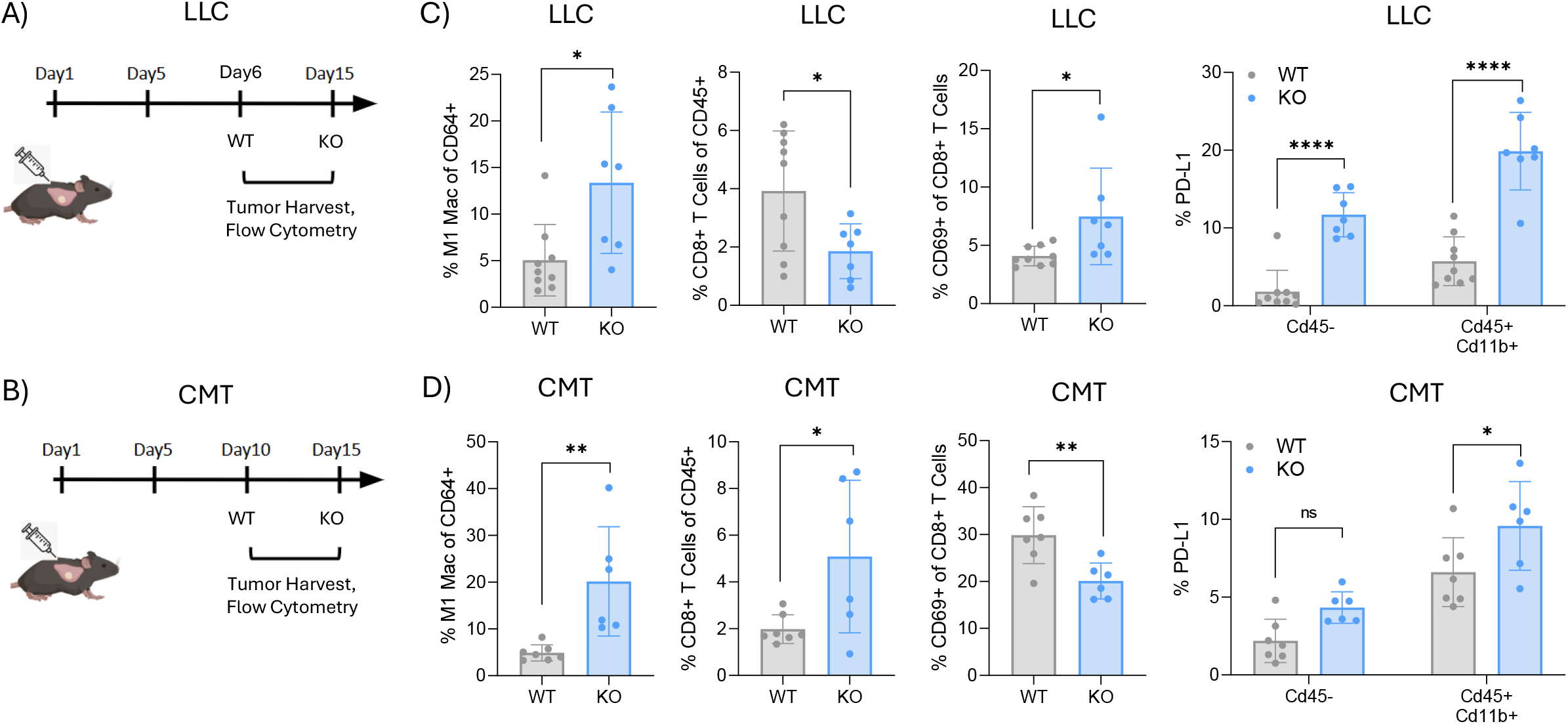
Increased prevalence of PD-L1+ cells in CD47-deficient tumours. Flow cytometry was used to conduct immune profiling on LLC and CMT tumours harvested at the time points indicated in **A** and **B. C)** Frequency of M1-type macrophages (Mac, CD64+MHC-II^high^CD80+), CD8+ T cells, CD69+ CD8+ activated T cells, PD-L1+ CD45-cells and PD-L1+ CD45+ CD11b+ myeloid cells within LLC tumours. **D)** The same immune cell populations in CMT tumours. Immune phenotyping was conducted on N=9 LLC WT, N=7 LLC CD47 KO, N=7 CMT WT, and N=6 CMT CD47 KO tumours. Cell frequencies were compared between genotypes using unpaired two-tailed t-tests. Error bars indicate mean ± standard deviation.

Given this result, we tested PD-L1 expression in CD47 KO cells and cells treated with anti-CD47 antibodies *in vitro*. We observed similar PD-L1 expression in WT and CD47 KO cells, and in WT cells treated with anti-CD47 antibodies or isotype control **(Fig S4A-B)**, suggesting upregulation of PD-L1 in CD47-deficient tumours is an *in vivo*-specific phenomenon. These observations indicate that CD47-deficiency promotes tumour immunity and suggest that CD47 KO tumours compensate for their fitness disadvantage by utilizing PD-L1 as an alternative immunosuppressive mechanism.

### 3.3 Evidence of mutually exclusive expression of CD47 and CD274 in lung tumours

Expression patterns of CD47 and PD-L1 measured using immunohistochemistry (IHC) have been described as negatively correlated in some NSCLC studies.^15,16^ This suggests lung tumours may express one or the other to facilitate tumour immune escape. To investigate this idea, we used cBioPortal^9–11^ to examine the expression of *CD47* and the gene encoding PD-L1 (*CD274*) in lung adenocarcinoma (LUAD) tissues from the TCGA cohort.^17^ This revealed that *CD274* and *CD47* were upregulated relative to non-malignant tissues in 10% (52/510) and 6% (33/510) of tumours, respectively, with only one tumour having expression of both (0.2%; 1/510) **(Fig 4)**. Similar to reported IHC studies, these RNA expression data indicate a trend towards mutual exclusivity of *CD274* and *CD47* in the TCGA LUAD cohort **(Fig 4)**.

**Figure 4.**
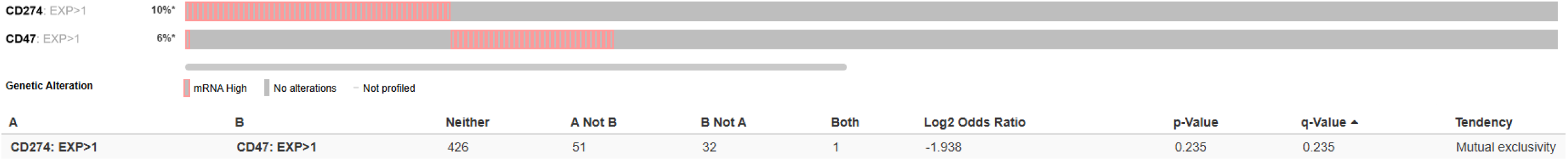
Trend of mutual exclusivity between CD47 and CD274 expression in TCGA lung adenocarcinoma. Upper, Oncoprint indicating RNA expression of *CD47* and *CD274* in 510 lung adenocarcinoma tumours from The Cancer Genome Atlas cohort. Z-scores for RNA expression were calculated relative to expression in normal tissue controls. A Z-score threshold > 1 was used to classify tumours as having high expression of *CD47* or *CD274*. Each column represents an individual tumour and pink indicates high mRNA expression. Lower, summary statistics for *CD47* and *CD274* expression in LUAD tumours indicating a trend towards mutual exclusivity. The plot was generated and mutual exclusivity analysis was done in cBioPortal.

### 3.4 PD-L1 blockade prolongs the survival of syngeneic mice with CD47-deficient tumours in the CMT model

Our findings suggest that PD-L1 may confer resistance to CD47 inhibition, providing a biological rationale for combining CD47- and PD-L1-targeted immunotherapies to mitigate acquired resistance and potentiate the efficacy of CD47 blockade. To test this, we further engineered our Cas9+ LLC and CMT WT and CD47 KO cells with or without PD-L1 LOF (sgCD274 and sgLacZ, respectively) **(Fig 5A)**. Then, we orthotopically injected WT control, single LOF (CD47 or PD-L1), or dual LOF (CD47 and PDL1) cells into syngeneic hosts. Mice were sacrificed at humane endpoints associated with tumour burden. These studies revealed that mice with CD47 KO + sgLacZ tumours displayed significantly longer survival times than mice with WT + sgLacZ tumours for both models, confirming our earlier results **(Fig 5B-C, Fig 2B,D)**. We observed model-specific responses to PD-L1 LOF alone and dual CD47 + PD-L1 LOF. For LLC, PD-L1 LOF in tumours conferred a significant survival benefit compared to WT controls. Dual LOF in LLC tumours did not provide an additional survival advantage compared to CD47 or PD-L1 LOF alone, although growth of tumours with dual LOF measured using BLI was initially impaired followed by rapid progression (**Fig 5B, Fig S5)**. In contrast, PD-L1 LOF in CMT tumour-bearing mice had no significant effect on survival time compared to WT controls, but mice bearing tumours with dual CD47 + PD-L1 LOF exhibited prolonged survival compared to mice with CD47 or PD-L1 LOF tumours alone **(Fig 5C)**.

**Figure 5.**
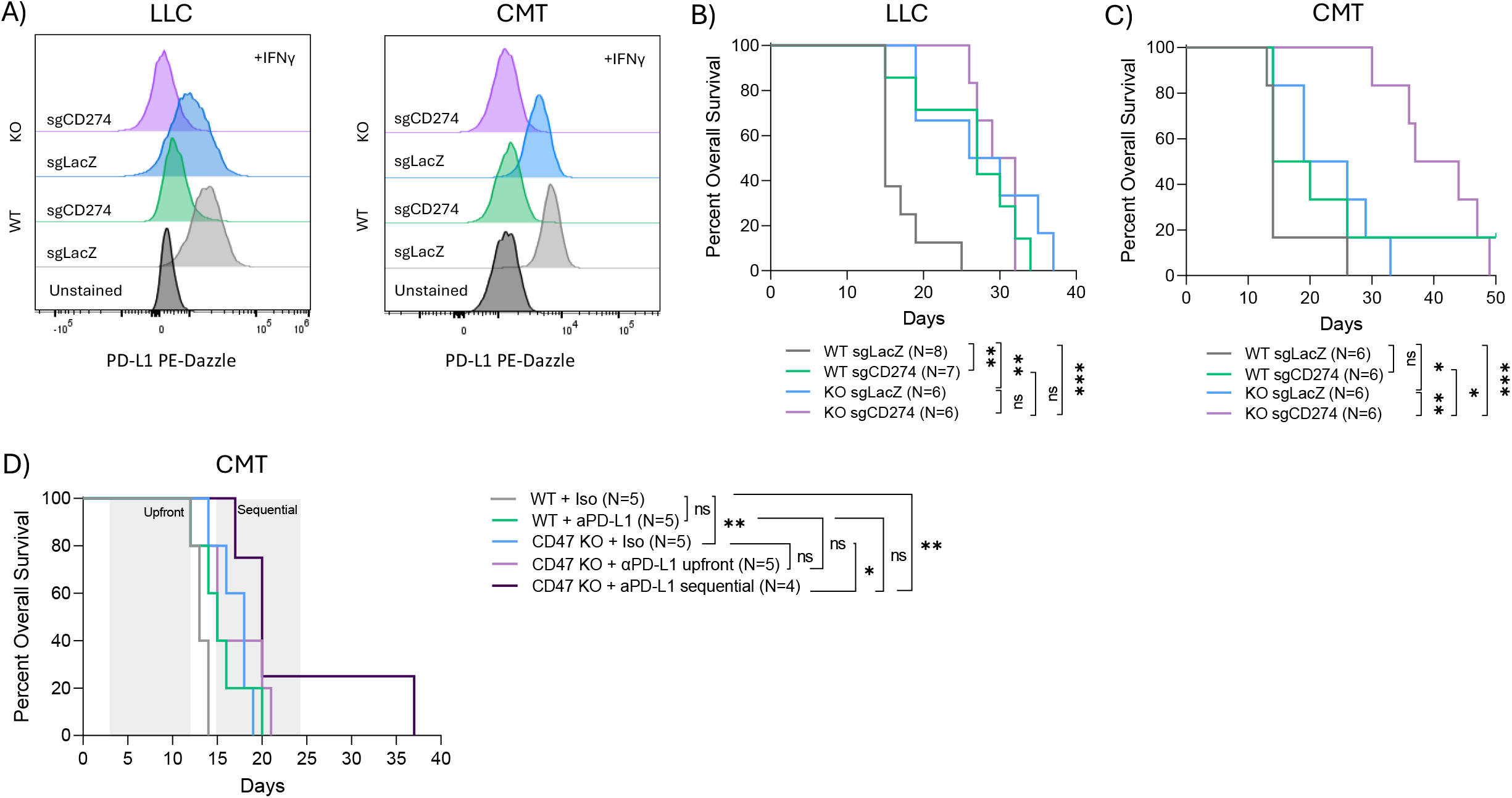
Combining CD47 and PDL1 blockade prolongs survival in syngeneic tumour-bearing mice. **A)** Flow cytometry verifying loss of PD-L1 protein expression in LLC and CMT WT and CD47 KO cells. All cells were treated with 100ng/ml IFNy for 24hr to induce PD-L1 expression. **B**,**C)** Kaplan-Meier survival curves for mice orthotopically injected with WT, PD-L1 LOF, CD47 LOF or dual PD-L1 + CD47 LOF cells. For LLC, N=6-8 mice/genotype and for CMT, N=6/genotype, as indicated. Asterisks indicate significance for comparing survival curves across genotypes using log-rank tests. **D)** Kaplan-Meier survival curves for mice orthotopically injected with CMT WT or CD47 KO tumours treated with isotype control or anti-PD-L1 therapeutic antibody. Four doses of 200µg anti-PD-L1 antibody (clone 10F.9G2) were administered by intraperitoneal injection, given “upfront” to model concurrent CD47 and PD-L1 blockade, or starting on Day 15 to model “sequential” treatment (N=4-5/group as indicated). Grey boxes indicate treatment periods. Asterisks indicate significance for comparison of survival curves across treatment arms using log-rank tests. Iso = isotype control (clone LTF-2).

To corroborate our findings using an orthogonal approach, we conducted pharmacologic studies. We selected the CMT model because PD-L1 LOF extended the survival of mice with CD47-deficient tumours, and unlike LLC, CMT exhibits sensitivity to PD-1/PD-L1 blockade administered with therapeutic antibodies **(Fig S6)**.^12,18^ Since pilot studies revealed that anti-CD47 therapeutic antibodies do not impair the growth of lung tumours like genetic KO does in syngeneic mice **(Fig S7)**, we used CD47 KO lines to model CD47 blockade. CMT WT or CD47 KO cells were implanted orthotopically into mice and treated with isotype control or anti-PD-L1 antibodies beginning on Day 3 or Day 15 post-injection to model “upfront” combination or “sequential” therapy, respectively **(Fig 5D)**. Consistent with our genetic studies, anti-PD-L1 treatment of WT tumours had no significant survival effect, while isotype control-treated mice with CD47 KO tumours had longer survival times than WT tumour-bearing mice. Upfront anti-PD-L1 treatment of CD47 KO tumours elicited a non-significant survival benefit compared to CD47 KO and anti-PD-L1 alone. In contrast, initiating anti-PD-L1 therapy at the timepoint at which we observed an increased prevalence of PD-L1+ cells in CD47 KO tumours (i.e., “sequential therapy”) prolonged the survival of mice with CD47-deficient tumours compared to the same tumours treated with isotype control **(Fig 5D)**. These results suggest that administering PD-L1 blockade after CD47 inhibition most effectively suppresses orthotopic CMT tumour growth compared to single-agent blockade and upfront combination therapy. As such, PD-L1-targeted adjuvant therapy could be an effective strategy for enhancing the efficacy of CD47 blockade in lung cancer patients.

## 4.0 Discussion

To our knowledge, this is the first study to investigate tumour response to CD47 blockade using orthotopic syngeneic lung tumour models, with the intention of discovering resistance mechanisms that could be targeted to potentiate its anti-tumour efficacy. Our immunoprofiling of lung tumours isogenic for CD47 yielded insights into how its inhibition influences the immune milieu to stimulate tumour immunity. It also revealed an increased abundance of PD-L1+ cells in CD47-deficient tumours compared to WT controls. These findings, along with the results of our genetic and pharmacologic studies testing the effects of combined CD47 and PD-L1 inhibition, are consistent with PD-L1 mediating an adaptive resistance mechanism that limits the efficacy of CD47-targeted monotherapy.

Discovery of this putative resistance mechanism could be translated to inform patient selection for CD47 blockade and/or design effective combination strategies for maximizing its efficacy. With respect to the latter, PD-L1-targeted immunotherapy is a tractable solution for mitigating resistance to CD47 blockade in the clinic. The utility of upfront combination therapy is supported by a number of preclinical studies that have demonstrated anti-tumour efficacy upon dual inhibition of CD47 and PD-L1/PD-1 in solid tumour models. For example, concurrent CD47 and PD-1/PD-L1 blockade using monoclonal antibodies, bispecific antibodies, fusion proteins, or nanoparticle approaches has been shown to enhance tumour killing in models of several tumour types, including breast cancer, colorectal cancer, pancreatic cancer, lymphoma, and NSCLC.^16,19–29^

Interestingly, clinical studies have reported that CD47 and PD-L1 expression are anti-correlated in NSCLC tissues.^15,16^ This observation, combined with our work, suggests that PD-L1 could serve as a biomarker to contraindicate CD47-targeted immunotherapy in certain patients. It also supports the idea that lung tumours differentially exploit CD47 and PD-L1 to evade tumour immunity, which could explain why “sequential” treatment of CD47 KO tumours with anti-PD-L1 therapy prolonged survival compared to earlier “upfront” administration in our study, which may be less effective prior to PD-L1 upregulation. We hypothesize that tumours compensate for enhanced tumour immunity induced by CD47 inhibition by upregulating PD-L1 to facilitate immune suppression; and consequently, that sequential treatment with PD-L1-targeted immunotherapy is optimal for potentiating CD47 blockade because it targets the acquired dependence of tumours on PD-L1.

Additional studies to validate the biological relevance and therapeutic potential of our findings are needed to address the limitations of our study. First, constitutive genetically-mediated inhibition of CD47 and PD-L1 in some of our *in vivo* experiments does not recapitulate the physiological and pharmacodynamic effects of treating established tumours with drugs like monoclonal antibodies in patients. For example, we observed an increase in CD45+CD11b+PD-L1+ immune cells in CD47-deficient tumours, which would not be inhibited by tumour-specific PD-L1 KO in our genetic studies. Our use of KO to inhibit CD47 in tumours reflects challenges associated with systemic administration of monoclonal antibodies targeting CD47 due to its ubiquitous expression.^30^ Indeed, we observed limited efficacy of anti-CD47 monoclonal antibodies against LLC and CMT tumours grown in syngeneic mice, which is likely attributable to the CD47 antigen sink that reduces the therapeutic load reaching the tumour. This highlights the need for continued research to develop alternative modes of delivering CD47 blockade in a tumour-focused manner to realize its therapeutic effects, and this is an active area of translational research. Second, our immune profiling was performed at a single timepoint, which does not capture the dynamic evolution of changes in the tumour-immune milieu associated with CD47 inhibition over time. As such, the time at which PD-L1+ cells start to accumulate following CD47 blockade remains to be determined. Longitudinal studies to monitor immune phenotypes at multiple timepoints could identify the optimal window for administering PD-L1 therapy and additional acquired resistance mechanisms to inform other combination strategies for maximizing CD47 blockade efficacy. Finally, our studies are limited to syngeneic murine NSCLC models. Validation in tumour samples from patients treated with CD47 blockade in clinical trials, and/or human tumour xenografts grown in mice with humanized immune systems are needed to confirm the clinical relevance of our findings. Nevertheless, the putative PD-L1-mediated adaptive resistance mechanism we have discovered in syngeneic lung tumour models is logical based on the biological processes underlying the cancer immunity cycle, which stimulated the development of CD47-PD-L1 dual-targeting therapies.^31^

Collectively, our findings establish CD47 as a functionally and therapeutically relevant immune checkpoint in NSCLC. Our *in vivo* studies provide proof-of-concept and biological rationale for using PD-L1 checkpoint inhibitors as an adjuvant treatment following CD47 blockade to circumvent resistance and enhance its therapeutic activity. Our work has yielded clinically-relevant insights needed to guide effective use of CD47 blockade in NSCLC, and justifies additional research to further develop CD47 inhibitors and define their clinical utility.

## Abbreviations

KO: knockout
CMT: CMT167
LLC: Lewis Lung Carcinoma
NSCLC: non-small cell lung cancer
IP: intraperitoneal
mAb: monoclonal antibody
WT: wildtype
LOF: loss of function
Iso: isotype control

## 5.0 Author Credit Contributions

Asa P. Y. Lau - Conceptualization, Methodology, Investigation, Data acquisition, Data curation, Formal analysis, Visualization, Writing - original draft, Writing - review & editing, Project Administration

Kristyna A. Gorospe - Investigation, Writing - review & editing

Kelsie L. Thu - Conceptualization, Methodology, Investigation, Data curation, Writing - original draft, Writing - review & editing, Supervision, Funding Acquisition, Project Administration

## 7.0 Acknowledgements

The authors are grateful to the Unity Health Toronto Vivarium staff for assistance with animal studies, as well as Flow Cytometry and Bioimaging core scientists, Dr. Monika Lodyga and Dr. Caterina di Ciano-Oliveira for technical assistance.

## 10.0 Supplementary Materials

The following supplementary materials accompany this article and are available in SupplementaryData.pdf

Figure S1. Flow cytometry gating for tumour immune phenotyping

Figure S2. Representative BLI images for orthotopic LLC-Luc tumours in syngeneic mice Figure S3. Additional immune phenotypes in WT and CD47-deficient LLC and CMT tumours

Figure S4. PD-L1 expression in WT and CD47 KO cells and WT cells treated with anti-CD47 antibody.

Figure S5. Growth curves for LLC tumours with single and dual CD47 and PD-L1 LOF. Figure S6. Efficacy of PD-1 and PD-L1 blockade in syngeneic lung tumour models

Figure S7. Efficacy of antibody-mediated CD47 blockade in syngeneic lung tumour models

Table S1. Single guide RNA (sgRNA) sequences for CRISPR-mediated genetic editing Table S2. Antibodies used for flow cytometry

