## Supplementary Material for "Upregulation of PD-L1 as a putative mechanism of resistance to CD47 inhibition in non-small cell lung cancer"

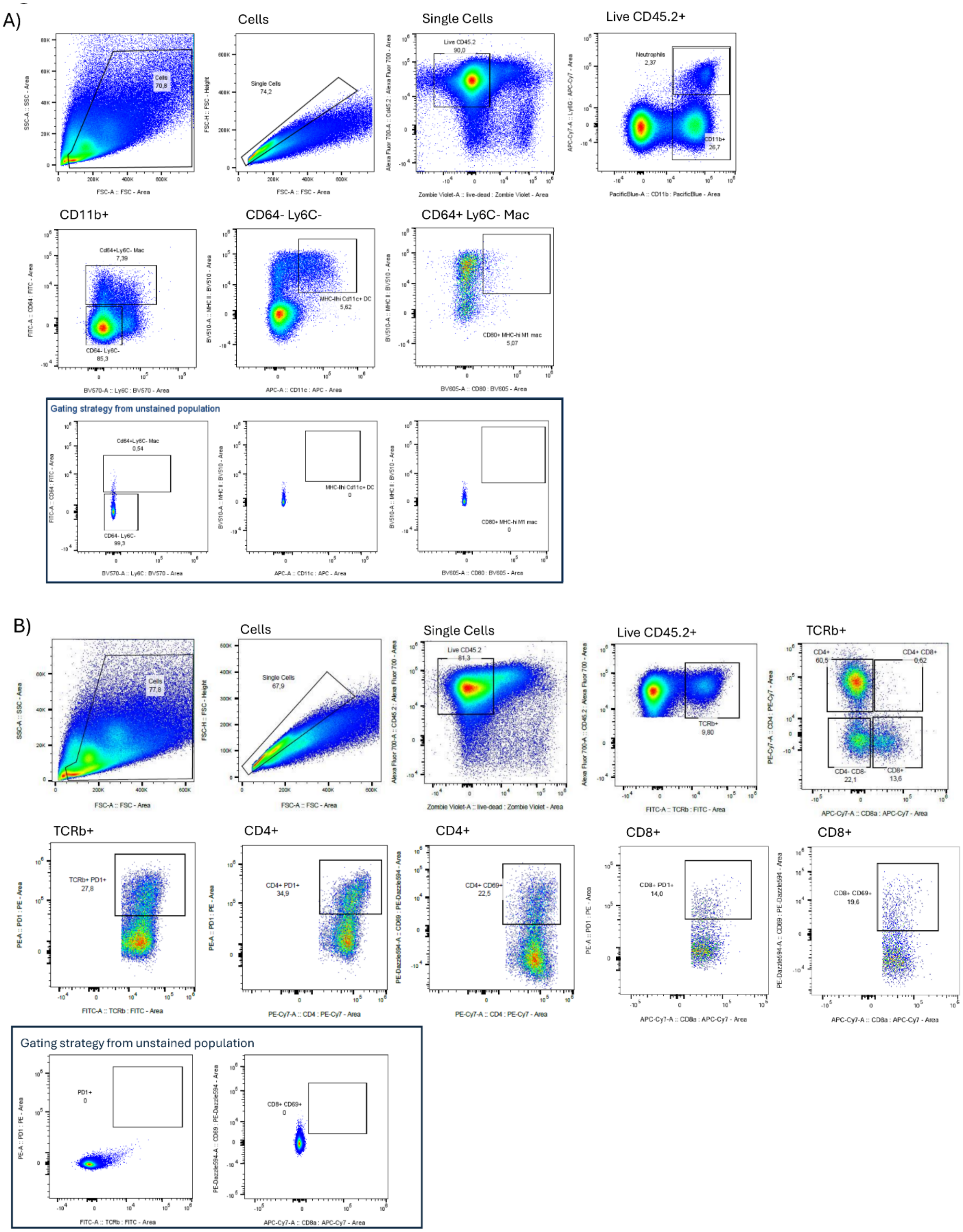
A)
Cells
Single Cells
Live CD45.2+
Cells 70,8
Single Cells 74,2
Live CD45.2 90,0
Neutrophils 2,37
CD11b+ 26,7
CD11b+
CD64- Ly6C-
CD64+ Ly6C- Mac
Cd64+Ly6C- Mac 7,39
CD64- Ly6C- 85,3
MHC-IIhi Cd11c+ DC 5,62
CD80+ MHC-hi M1 mac 5,07
Gating strategy from unstained population
Cd64+Ly6C- Mac 0,54
CD64- Ly6C- 99,3
MHC-IIhi Cd11c+ DC 0
CD80+ MHC-hi M1 mac 0
B)
Cells
Single Cells
Live CD45.2+
TCRb+
Cells 77,8
Single Cells 67,9
Live CD45.2 81,3
TCRb+ 9,80
CD4+ 60,5
CD4+ CD8+ 0,62
CD4- CD8- 22,1
CD8+ 13,6
TCRb+
CD4+
CD4+
CD8+
CD8+
TCRb+ PD1+ 27,8
CD4+ PD1+ 34,9
CD4+ CD69+ 22,5
CD8+ PD1+ 14,0
CD8+ CD69+ 19,6
Gating strategy from unstained population
PD1+ 0
CD8+ CD69+ 0

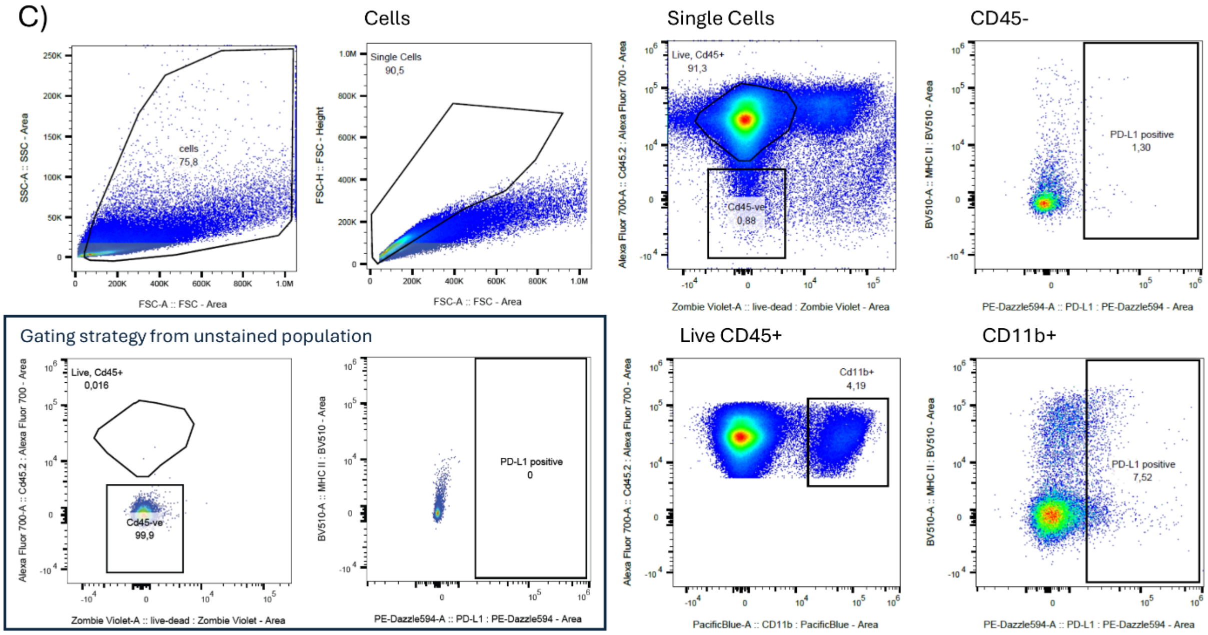


**Figure S1. Flow cytometry gating for tumour immune phenotyping. A)** Gating strategy for myeloid cells. Cells were gating on singlets, followed by live/dead discrimination. Live CD45.2+ cells were gated for CD11b+ myeloid cells and CD11b+ Ly6G+ neutrophils. CD11b+ cells were further gated based on CD64 and Ly6C expression. Dendritic cells were defined as CD64- Ly6C- Mhc-II$^{high}$ CD11c+. M1-polarized macrophages were defined as CD64+ Mhc-II$^{high}$ CD80+. **B)** Gating strategy for T cells. After gating for singlets and viable cells, live CD45.2+ cells were gated for TCRb+ T cells. CD4+ helper and CD8+ cytotoxic T cell subsets were identified within the TCRb+ population. PD-1+ and activated CD69+ T cell subsets were further defined within the CD4+ and CD8+ populations. **C)** Gating strategy for detecting PD-L1+ cells. Tumour cells were identified as CD45- and myeloid cells as CD45+ CD11b+. PD-L1+ cells were defined as the percentage within each population. All gates were set on unstained cells. All plots presented are for representative WT tumour samples.

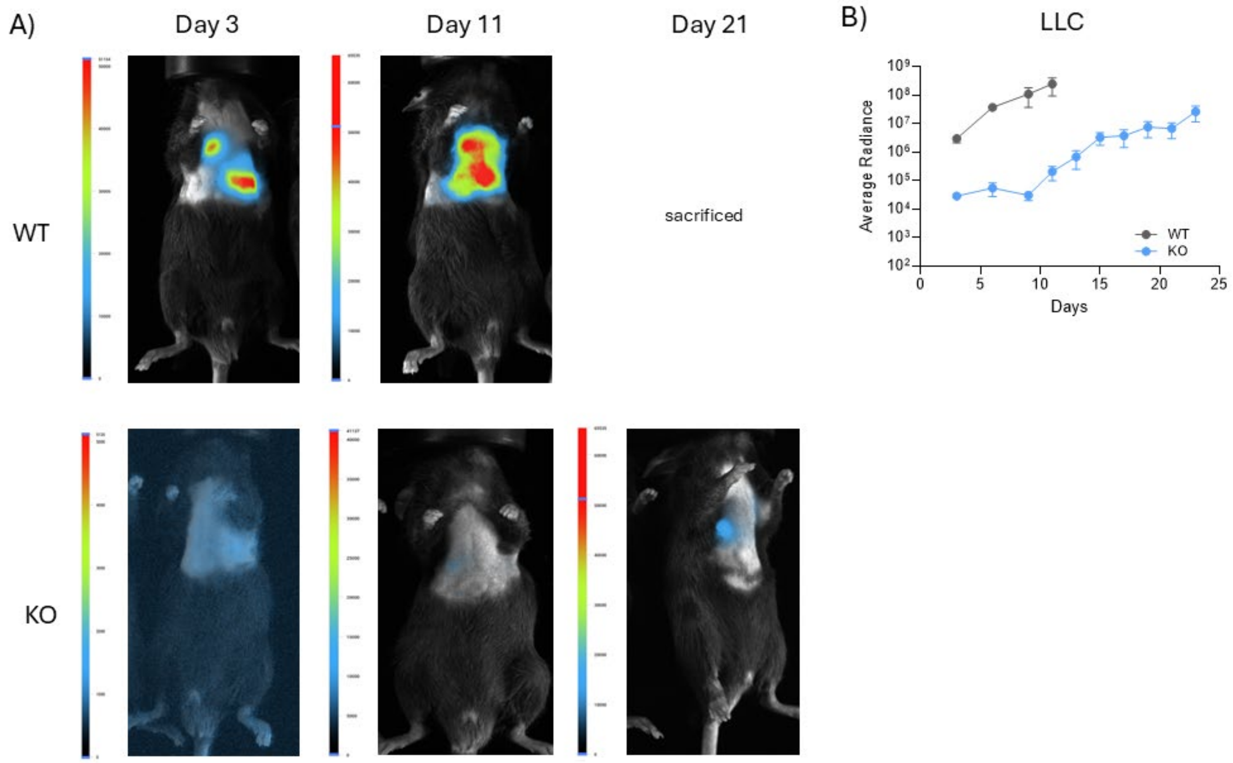


**Fig S2. Representative BLI images for orthotopic LLC-Luc tumours in syngeneic mice. A)** C57BL/6 mice were orthotopically implanted with LLC-Luc cells. BLI was performed as described in the methods to monitor tumour formation and growth from Day 3 onwards Representative mice with WT and CD47 KO tumours are shown. **B)** Curves were generated using bioluminescence imaging (BLI) to detect luciferase activity. Average radiance (photons/s/cm$^2$/sr) across replicate mice (N=6 for WT; N=7 for CD47 KO) is plotted. Error bars indicate mean ± SEM.

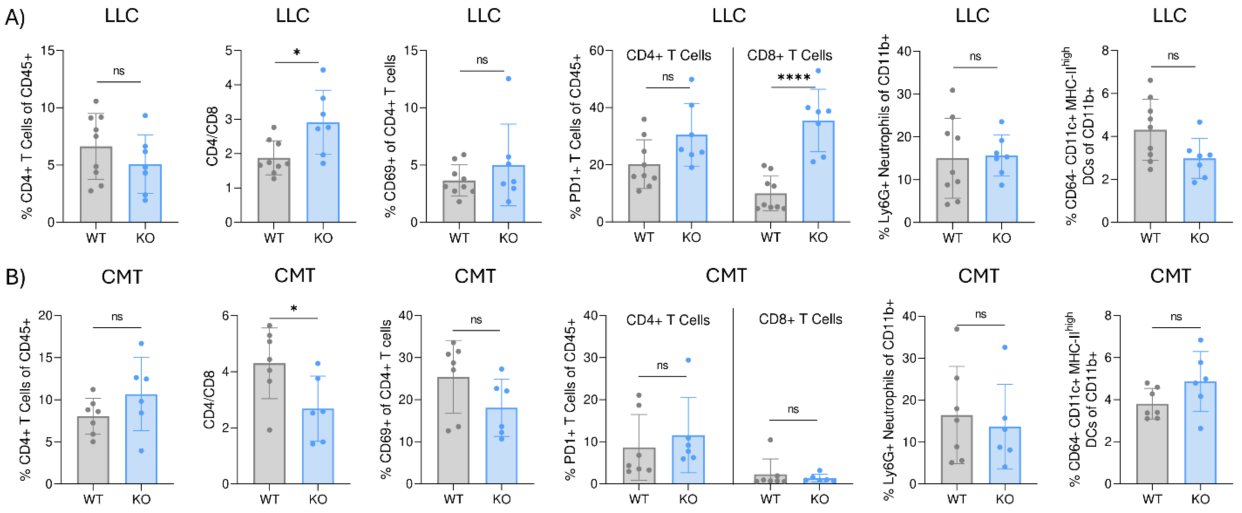


**Figure S3. Additional immune phenotypes in WT and CD47-deficient LLC and CMT tumours**. **A)** Ratio of CD4+/CD8+ T cells, frequency of CD4+ helper T cells, CD69+ CD4+ activated T cells, PD-1+ CD4+ and PD-1+ CD8+ T cells, neutrophils (CD11b+ Ly6G+) and dendritic cells (CD11b+ CD64- Ly6C- Mhc-II$^{hgh\ i}$CD11c+) within LLC tumours (WT N=9, KO N=7). **B)** The same immune cell populations in CMT tumours (WT N=7, KO N=6). Cell frequencies were compared between genotypes using unpaired two-tailed t-tests. Error bars indicate mean ± standard deviation.

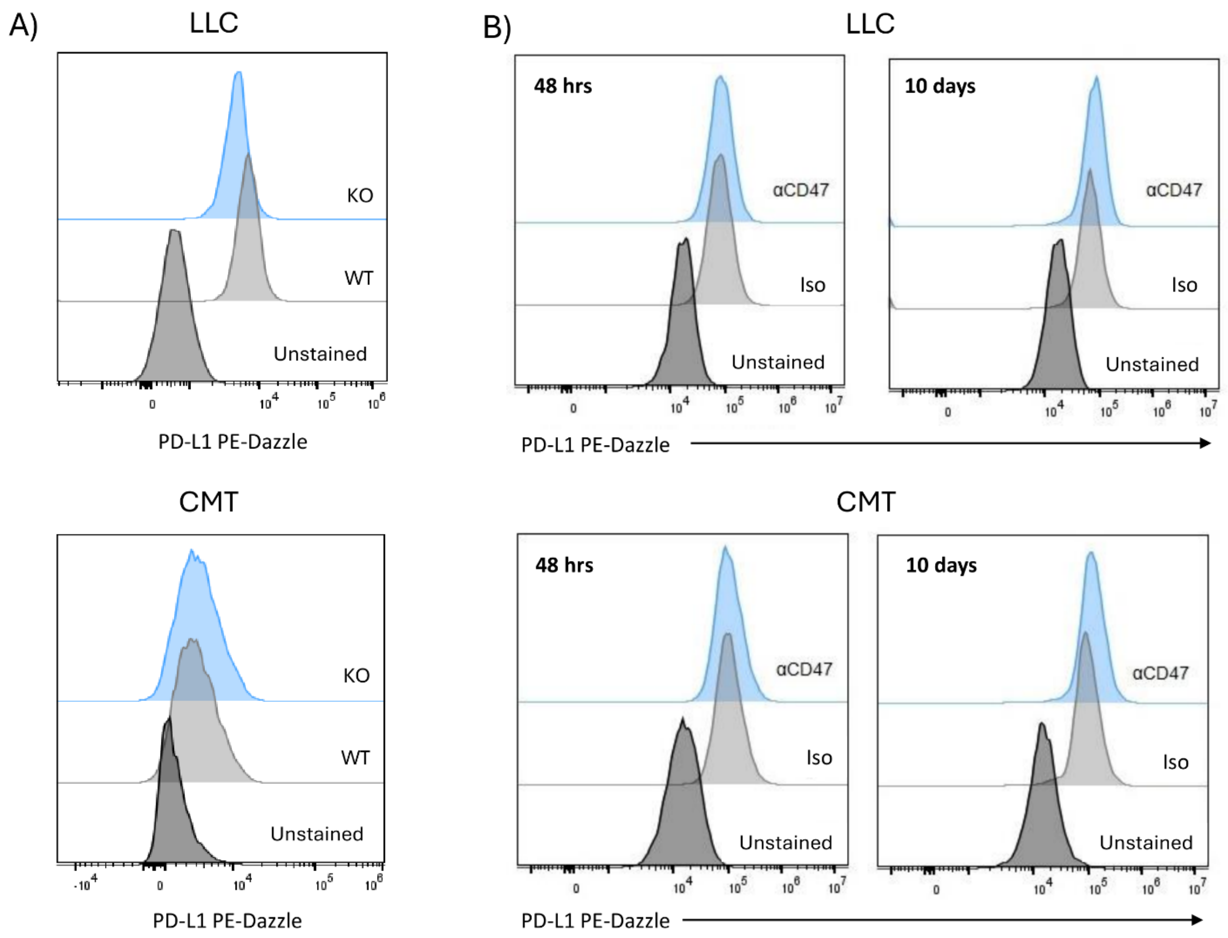


**Figure S4. PD-L1 expression in WT and CD47 KO cells and WT cells treated with anti-CD47 antibody.**

**A)** Surface expression of PD-L1 in WT and CD47 KO cells. **B)** Expression of PD-L1 in LLC and CMT cells treated with 0.01 mg/mL anti-CD47 antibody (αCD47; clone miap301) or rat isotype control (iso; clone 2A3) for 48 hours or 10 days. Expression was measured using flow cytometry. Representative plots are shown from 3 independent experiments.

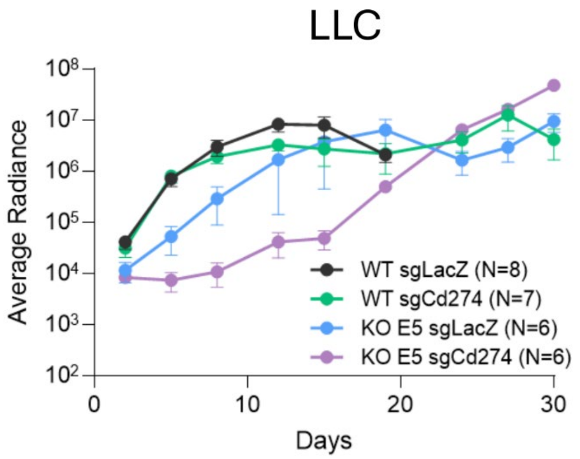


**Figure S5. Growth curves for LLC tumours with single and dual CD47 and PD-L1 LOF.** Growth curves were generated from BLI imaging data. Average radiance (photons/s/$cm^2$/sr) across replicate mice (N=6-8, as indicated) is plotted. Error bars indicate mean ± SEM.

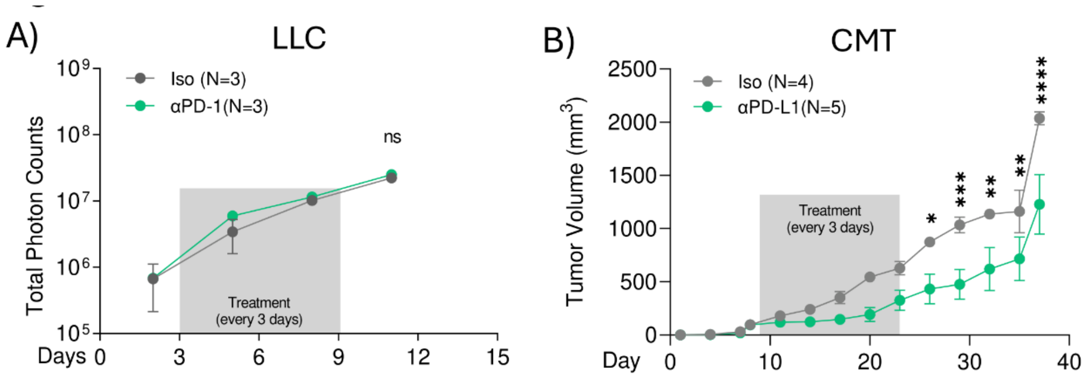


**Figure S6. Efficacy of PD-1 and PD-L1 blockade in syngeneic lung tumour models**. **A)** Orthotopic LLC and **B)** subcutaneous CMT tumours were treated with anti-PD-1 (clone RMP1-14; 150µg IP) or anti-PD-L1 (clone 10F.9G2 100µg IT), respectively, every 3 days. Grey boxes indicate treatment periods. Growth was monitored with BLI (LLC) or calipers (CMT). Asterisks indicate significance for two-tailed, unpaired t-tests. Error bars indicate mean ± SEM.

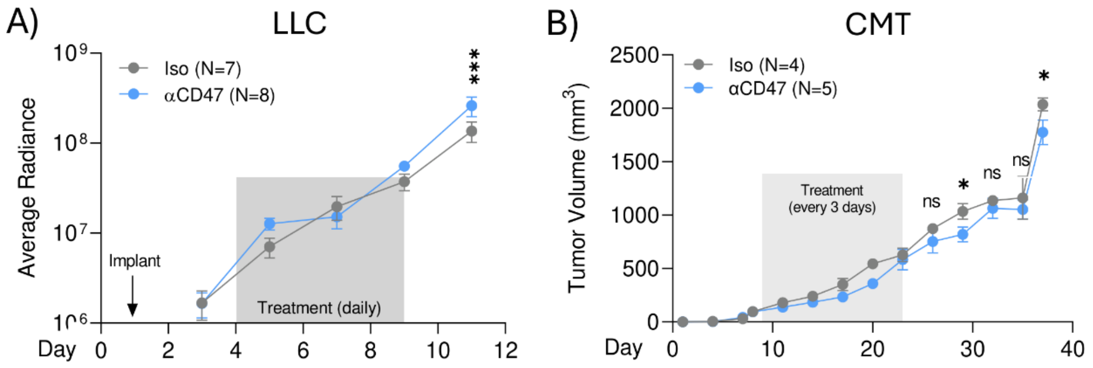


**Figure S7. Efficacy of antibody-mediated CD47 blockade in syngeneic lung tumour models**. **A)** Response of orthotopic LLC tumours treated with 200µg αCD47 (clone miap301; N=8) or isotype control (Iso; clone 2A3; N=7) delivered IP daily. Tumour growth was monitored with BLI. Radiance indicates ph/s/cm$^2$/sr. **B)** Response of subcutaneous CMT tumours treated with 100µg αCD47 (clone miap301; N=5) or isotype control (Iso; clone 2A3; N=4) delivered by intratumoural injection every 3 days. Tumour growth was monitored using calipers. For both models, grey boxes indicate the treatment period and error bars indicate mean ± SEM.

### Supplementary Tables

**Table S1. Single guide RNA (sgRNA) sequences for CRISPR-mediated genetic editing**

| Oligo | Guide Sequence |
|---|---|
| sgCD47-1 | TATAGAGCTGAAAAACCGCA |
| sgCD47-2 | CCACATTACGGACGATGCAA |
| sgCD274 | GCCTGCTGTCACTTGCTACG |

**Table S2. Antibodies used for flow cytometry**

| Marker | Fluor | Catalog Number (Biolegend) | Dilution |
|---|---|---|---|
| CD45.2 | AF700 | 109822 | 1:200 |
| TCRb | FITC | 109206 | 1:200 |
| CD8a | APC-Cy7 | 100714 | 1:200 |
| CD4 | PE-Cy7 | 100528 | 1:200 |
| CD69 | PE-dazzle | 104535 | 1:200 |
| CD137/4-1BB | APC | 106109 | 1:200 |
| CD279/PD-1 | PE | 135205 | 1:200 |
| CD11b | Pacific Blue | 101224 | 1:800 |
| CD11c | APC | 117310 | 1:200 |
| CD64 | FITC | 139316 | 1:200 |
| Ly6C | BV570 | 128017 | 1:500 |
| Ly6G | APC-Cy7 | 127623 | 1:200 |
| MHC-II | BV510 | 107639 | 1:750 |
| CD172a (SIRPa) | PE | 144011 | 1:200 |
| CD274 (PD-L1) | PE-dazzle | 124323 | 1:200 |
| F4/80 | PerCP-Cy5.5 | 123107 | 1:200 |
| CD80 | BV605 | 104729 | 1:100 |
| CD47 | APC | 127514 | 1:100 |
| CD47 | AF647 | 127509 | 1:200 |
